# Choice of 3D morphometric method leads to diverging interpretations of form–function relationships in the carnivoran calcaneus

**DOI:** 10.1101/2022.05.16.492149

**Authors:** Alexa N. Wimberly, Rossy Natale, Robert Higgins, Graham J. Slater

**Affiliations:** Department of Organismal Biology and Anatomy, University of Chicago; Department of the Geophysical Sciences, University of Chicago; Department of Earth Sciences University College London, U.K.

**Keywords:** Locomotion, Allometry, Macroevolution, Ecomorphology, Mammal

## Abstract

Three dimensional morphometric methods are a powerful tool for comparative analysis of shape. However, morphological shape is often represented using landmarks selected by the user to describe features of perceived importance, and this may lead to over confident prediction of form-function relationships in subsequent analyses. We used Generalized Procrustes Analysis (GPA) of 13 homologous 3D landmarks and spherical harmonics (SPHARM) analysis, a homology-free method that describes the entire shape of a closed surface, to quantify the shape of the calcaneus, a landmark poor structure that is important in hind-limb mechanics, for 111 carnivoran species spanning 12 of 13 terrestrial families. Both approaches document qualitatively similar patterns of shape variation, including a dominant continuum from short/stout to long/narrow calcanea. However, while phylogenetic generalized linear models indicate that locomotor mode best explains shape from the GPA, the same analyses find that shape described by *SPHARM* is best predicted by foot posture and body mass without a role for locomotor mode, though effect sizes for all are small. User choices regarding morphometric methods can dramatically impact macroevolutionary interpretations of shape change in a single structure, an outcome that is likely exacerbated when readily landmarkable features are few.

## Introduction

Skeletal morphology provides a crucial source of data for understanding evolutionary processes over time, space, and phylogeny. Functional demands on the skeleton related to behaviors such as feeding and locomotion frequently lead to predictable relationships between an organism’s morphology and its ecology (Wainwright, 1991; Bock, 1994; Barr, 2018). In turn, these form–function relationships allow for the inference of behavior in species for which we have only morphological data, such as fossils (Chen and Wilson, 2015; Nations et al., 2019; Grossnickle et al., 2020; Lungmus and Angielczyk, 2021), quantification of macroevolutionary rates and modes (Kilbourne and Hutchinson, 2019; Law et al., 2019; Law, 2021; Prang et al., 2021; Slater, 2022), and the testing of hypotheses about ecological responses to competition and environmental change (Feder et al., 2010; Polly, 2010; Polly et al., 2017; Short and Lawing, 2021). A number of approaches have been used to quantify patterns of ecomorphological variation in the post-cranial skeleton, ranging from functional indices derived from linear measurements (Van Valkenburgh, 1987; Losos, 1990; Garland and Janis, 1993; Collar et al., 2013; Barr, 2014) to the description of complex patterns of 3D shape variation using the tools of geometric morphometrics (Curran, 2012; Fabre et al., 2013; Martín-Serra et al., 2014; Wang et al., 2020; Dunn and Avery, 2021). However, the degree to which these different approaches to the quantification of organismal form may yield different ecological interpretations remains largely unexplored (but see Gould, 2014).

The calcaneus, or heel bone, has emerged as one of the most useful elements in the mammalian postcranial skeleton for the study of form, function, ecology, and locomotion (Polly, 2010; Polly and Sarwar, 2014; Polly et al., 2017; Panciroli et al., 2017; Polly, 2020). The calcaneus of therian mammals is made up of a distal head and an elongate proximal tuber (figure 1). Much of the calcaneus articulates distally with the cuboid, and with the astragalus in various orientations in different mammalian groups. The proximal end of the elongate tuber of therian mammals serves as the attachment site for the gastrocnemius and soleus muscles via the Achilles tendon, and serves as the in-lever for plantar flexion. A laterally placed structure, which shows various shapes in different mammal groups, is a tubercle-like process for peroneal muscle function in therian mammals; in carnivorans the peroneal process shows a groove for the tendon of the peroneus brevis and peroneus longus muscles to pass over to distal tarsals and metatarsals, essential for eversion of the foot (Polly, 2008). Measurements and shape studies on the calcaneus have yielded important insights into taxonomic identification (Stains, 1959, 1962), prediction of body mass (Yapuncich et al., 2015; Tsubamoto, 2019), inference of paleoenvironments (Youlatos, 2003; Kovarovic and Andrews, 2007; Curran, 2012, 2015), and even assessment of individual maturity (Jogahara and Natori, 2013). Because of its robust nature, the calcaneus is well-represented in the mammalian fossil record and its morphology has been used for predicting locomotor mode in fossils from disparate mammal clades, such as artiodactyls (Kovarovic, 2004; Curran, 2015; Kovarovic and Andrews, 2007), marsupials (Bassarova et al., 2009), primates (Boyer et al., 2013), and xenarthrans (Jasinski and Wallace, 2014).

**Figure 1:**
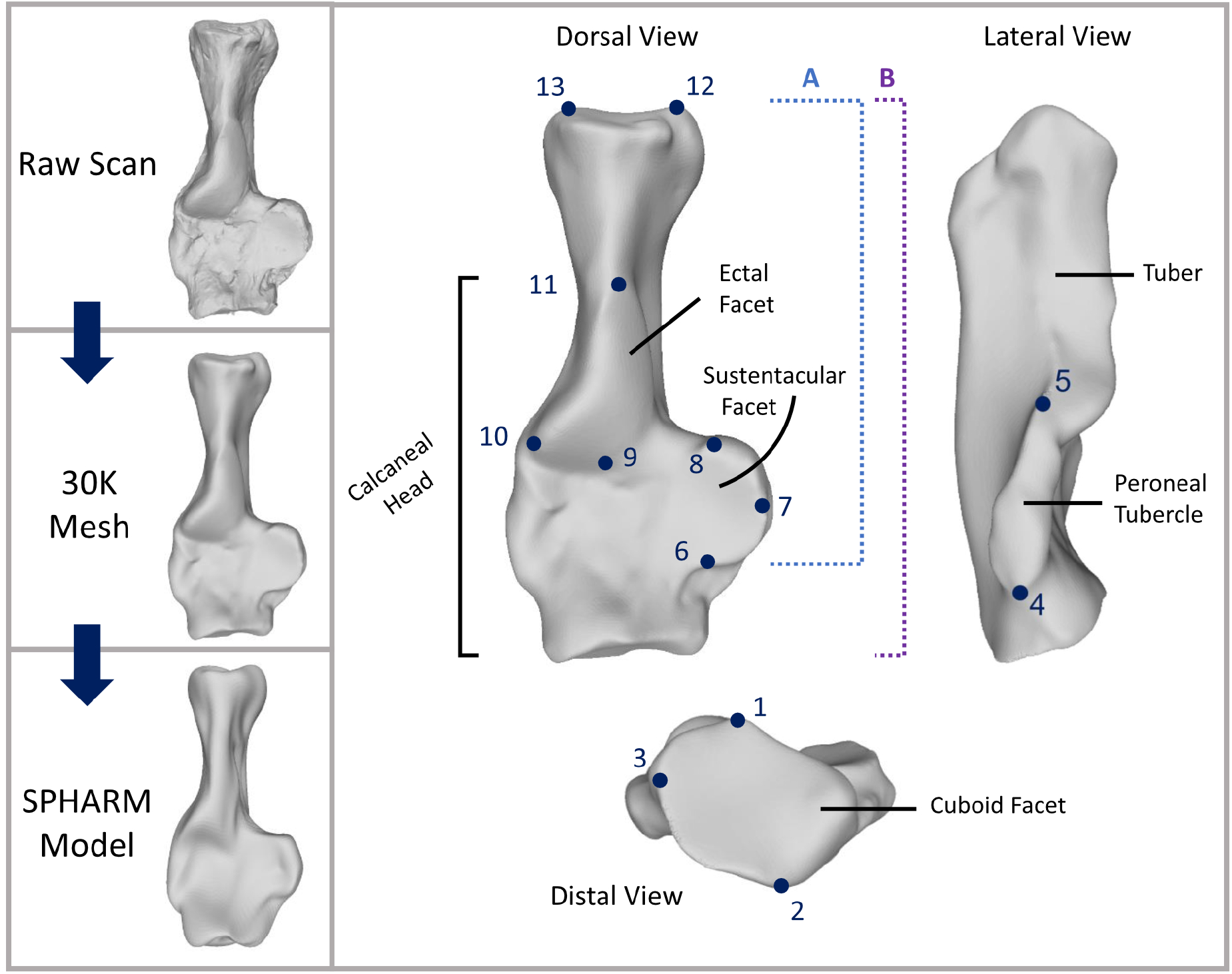
(Left) Figure showing the processing steps for our spherical harmonics analysis starting with a raw scan, decimating to 30,000 triangles, then generating the *SPHARM* model. The calcaneus represented here is that of *Nandinia binotata*, the African palm civet, which we chose as our template for spherical harmonics. (Right) Labeled anatomy of a right calcaneus (*Nandinia binotata*) with numbered landmarks. The landmarks are described as (1) dorsalmost point of the cuboid facet, (2) ventralmost point of the cuboid facet, (3) lateralmost point of the cuboid facet, (4) distalmost point of the peroneal tubercle, (5) proximalmost point of the peroneal tubercle, (6) distalmost point of the sustentacular facet, (7) medialmost point of the sustentacular facet, (8) proximalmost point of the sustentacular facet, (9) medio-distal most corner of the ectal (or calca-neoastragalar) facet, (10) lateralmost point of the ectal facet, (11) proximal end of the ectal facet, medial tip of the proximal tuber, and (13) the lateral tip of the proximal tuber. The dotted blue line (A) represents the length from the proximal tip of the tuber to the distal-most part of the sustentaculum, while the dotted purple line (B) represents the entire calcaneus length. Gear ratio is measured as B/A, or full calcaneal length divided by sustentacular length.

Calcaneal shape variation has been particularly intensely studied in Carnivora, an order of 304 extant species with extremely diverse diets, habitats, body masses, and locomotor modes. Carnivorans exhibit numerous postcranial specializations that allow some species to achieve hindfoot reversal (Jenkins Jr and McClearn, 1984; Taylor, 1988; Morales et al., 2018), swimming proficiency (Polly, 2008; Botton-Divet et al., 2017, 2018), and incredible running speeds (Hudson et al., 2012). Researchers have applied a number of two- and three-dimensional methods to quantify the shape of the calcaneus in carnivorans and assess its relationship with locomotor behavior, body mass, and habitat. A simple ecomorphological proxy is the calcaneal gear ratio (Polly, 2010; Polly and Sarwar, 2014; Polly et al., 2017; Polly, 2020). Gear ratio, which is the length of the out lever of the gastrocnemius and soleus muscles (full length of the calcaneus) divided by the length of their in lever (sustentacular length; fig. 1), serves as a proxy for hindlimb mechanics and is highly correlated with foot posture (Polly, 2010), locomotion, phylogeny, and habitat (Polly and Sarwar, 2014; Polly et al., 2017; Polly, 2020). Panciroli et al. (2017) explored a more comprehensive set of linear measurements of extant carnivoran calcaena, as well as landmarks digitized on photographs of calcanea in dorsal view, and used discriminant analyses and multivariate analyses of variance to find that about half the variation in calcaneal shape could be attributed to differences in locomotor mode, especially in ambulatory (e.g., ursids) and arboreal species.

A few studies have investigated the three-dimensional shape of the carnivoran calcaneus, but these have been phylogenetically and geographically restricted in sampling. Polly (2008) used an interpolation approach to spread an equal number of landmarks on a 3D “fishnet” point grid on surface scans of the calcanea and astragali of 12 extant carnivoran species, finding no relationship between body mass and multivariate shape, but varying strengths of relationships for locomotor mode, foot posture, and number of toes. Using a refined version of this approach, Polly and MacLeod (2008) attempted to classify extinct carnivorans to locomotor mode, foot posture, and digit number classes, with mixed success. Polly et al. (2017) explored 3D calcaneal shape in a larger sample of 37 North American carnivorans, finding high phylogenetic signal. However, their data comprised only 13 homologous landmarks representing characteristics associated with articular surfaces and did not explore the relationship between multivariate shape and locomotor behavior, size, or foot posture. As a result, it remains to be seen whether the ecomorphological signals that emerge in low dimensional studies of carnivoran calcaneal shape are maintained when considering full 3D shape across the entire order.

Here, we quantify the shape variation in 111 carnivoran calcanea spanning 12 of 13 terrestrial families using two distinct three-dimensional methods to answer questions about locomotor and ecological diversity. We first use standard geometric morphometric methods to quantify shape variation based on thirteen 3D landmarks (Polly et al., 2017). Next, we use spherical harmonics (*SPHARM*: McPeek et al., 2008; Shen et al., 2009) as a more holistic approach to quantifying the full 3D shape of the calcaneus. Using phylogenetically informed comparative methods, we then evaluate phylogenetic and functional signals in carnivoran calcaneal shape from the two datasets. Our results reveal that the choice of three-dimensional method can significantly alter interpretation of form–function relationships.

## Materials and methods

### 3D Scanning

Carnivoran specimens used in this study were acquired from the mammal collections of The Field Museum of Natural History (Chicago, IL). We sampled one right calcaneus when possible from 111 of terrestrial carnivores that span 12 of 13 terrestrial families; calcanea of *Prionodon* (Prionodontidae) were not available. If the right calcaneus was in poor condition or unavailable, we scanned the left calcaneus and digitally mirrored the specimen. We only used adult calcanea, as indicated by the fully fused growth plate on the proximal part of the calcaneal tuber.

Surface scans of each specimen were generated using one of three structured light scanners depending on the size of the calcaneus. Small specimens were scanned using a Capture Mini blue light scanner (3D Systems, Rock Hill SC) at a resolution of 80 *µ*m or a Comet L3D blue light scanner (Zeiss, Jena, Germany) with 45 (0.18 *µ*m point spacing) or 75mm (40 *µ*m point spacing) field of view lenses. Larger specimens were scanned using a Go!Scan20 handheld white light scanner (Creaform, Levis Canada) at 0.2 mm resolution. We used Geomagic DesignX (3D Systems, Rock Hill SC) to clean and smooth models by removing floating polygons and correcting artificial blemishes in the model. The meshes were then refined by subdividing to increase the number of triangles, smoothing the entire model using the highest setting, re-meshing to create a solid, watertight model, and then decimating to 30,000 triangles. After ensuring that each decimated model was clean, watertight, and had no floating or manifold vertices, we exported them in stereolithographic (.STL) format.

### Ecological Data

We collected data on three putative predictors of calcaneal shape: body mass, locomotor mode, and foot posture. Estimates of average adult body mass were collected from the PanTHERIA database (Jones et al., 2009, Table S1). Categorical data for locomotor mode and foot posture were collected from the literature (Table S2,3)(Fig. 2). We classified species as terrestrial, scansorial, arboreal, natatorial or semi-fossorial based on how much they use a substrate for important activities such as resting, foraging/hunting, hiding, and traveling (Eisenberg, 1981). Terrestrial species are those that do not actively use other substrates for important activities (e.g., the coyote *Canis latrans* and lion *Panthera leo*). Species were considered scansorial if they were otherwise terrestrial, but use trees for significant purposes such as resting, foraging, or avoiding competing species and/or predators (e.g., the jaguar *Panthera onca* and coatimundi *Nasua nasua*). Species that extensively use trees for resting, foraging, mating, traveling, or evasion were considered arboreal (e.g., kinkajou *Potos flavus* and binturong *Arctictis binturong*). Species were assigned to the semi-fossorial group if they rely heavily on digging for shelter and/or foraging (e.g., the honey badger *Mellivora capensis* and spotted skunk *Spilogale putorius*). Finally, if water is ecologically significant for feeding or traveling, species were assigned to the natatorial category (e.g., otters and the marsh mongoose *Atilax paludinosus*). Species considered in previous studies to be “cursorial” were classified in our dataset as terrestrial; although several taxa exhibit adaptations for high speeds or pursuit predation over long distances (e.g. the cheetah *Acinonyx jubatus* and African wild dogs *Lycaon pictus*), these behaviors are frequently defined based on morphology (Carrano, 1999) leading to potential circularity. Ursids, in previous studies, have been labeled as “ambulatory” (e.g., Panciroli et al., 2017). We consider ambulatory to represent the union of terrestrial or scansorial locomotion, plantigrade foot posture, and large body size and therefore do not consider it to be a unique locomotor category.

**Figure 2:**
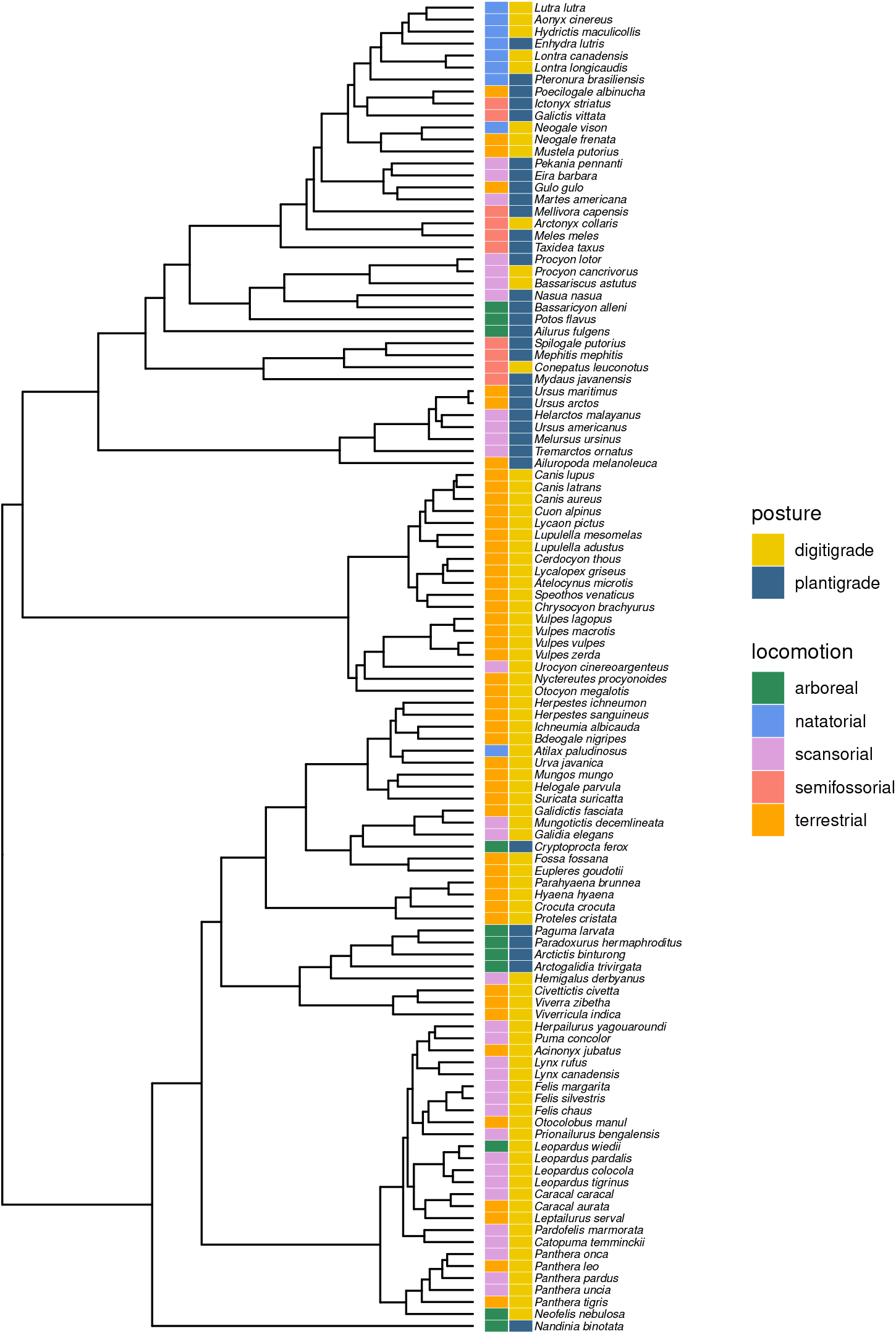
Phylogenetic tree of the 111 terrestrial carnivoran species included in this study. Two columns of colored boxes indicate (left) locomotor mode and (right) foot posture.

Data on foot posture were collected from previous studies of carnivorans (Carrano, 1997; Polly, 2008; Polly and MacLeod, 2008; Polly et al., 2017) and supplemented with other literature sources if no data were available or there were conflicting assignments (Table S3). Our foot posture categories are divided into plantigrade, where the entire sole of the foot, including the heel, contacts the substrate, and digitigrade, where only the digits consistently contact the ground while moving (Fig. 2). We omitted the use of semi-digitigrade because the degree of separation between the heel and the ground is extremely variable for this category. Thus, we assigned previously considered semi-digitigrade taxa as digitigrade.

### 3D shape analysis

We quantified 3D shape of carnivoran calcanea using two approaches. We first used Generalized Procrustes Analysis (GPA: Gower, 1975; Rohlf and Slice, 1990) to generate 3D shape coordinates from 13 3D landmarks (Fig. 1: Polly et al., 2017) digitized on each calcaneus model using `MeshLab` v2016.12 (Cignoni et al., 2008). Landmark files were read into `R` (R Core Team, 2021) using a modified version of a script by Randi Griffin (http://www.randigriffin.com/2017/05/07/read-meshlab-pickedpoints.html) and subjected to GPA to remove the effects of scale, translation, and rotation using the `gpagen` function in the `geomorph` library (Baken et al., 2021; Adams et al., 2021). The resulting aligned Procrustes coordinates were then projected into the linear tangent space to yield Procrustes shape variables that are suitable for statistical analysis. We use the function `gm`.`prcomp` to perform a principal components analysis on the Procrustes shape variables and visualized shape change along the axes of maximal shape variation using the function `plotRefToTarget`, which compares shapes corresponding to maximal scores along individual principal components to a reference shape using thin plate splines. We used the mean shape, obtained via the `mshape` function, as our reference shape.

GPA methods typically use landmark coordinates placed at homologous points on a surface or outline. As a result, these approaches may not fully describe the shape variation among morphological structures that exhibit complex patterns of shape variation but have few obvious landmarks, such as mammalian calcanea. To generate a more complete description of shape variation in carnivoran calceanea, we used spherical harmonic analysis (*SPHARM*). *SPHARM* is a 3D form of Fourier analysis that allows for examination of complex shape in a homology-free environment (McPeek et al., 2008; Shen et al., 2009). As such, spherical harmonic modeling has proven useful in quantifying axes of shape variation in structures that lack numerous homologous landmarks, such as damselfly reproductive structures (McPeek et al., 2008, 2009), carnivoran frontal sinuses (Curtis and Van Valkenburgh, 2014), and mammalian cervical vertebrae (Vander Linden et al., 2019). Spherical harmonic analyses were carried out using *SPHARM* v.1.5, through `MatLab` v 2017b (MathWorks Inc, Natick, Massachusetts). Models were first resized to unit centroid size. *SPHARM* analyses require all models to be aligned to a reference specimen through a minimal set of landmarks. We used the African palm civet *Nandinia binotata* as our template due to its generalized morphology. Specimens were aligned to the reference model using the thirteen landmarks digitized for GPA (Fig. 1). The aligned and scaled models were then subjected to iterative smoothing and used to construct spherical parameterizations. *SPHARM* models converge on a more accurate representation of the original object as the degree of the model is increased Shen et al. (2009). However, the number of spherical harmonic coefficients increases as the square of the degree plus 1 McPeek et al. (2008), resulting in larger and more computationally challenging datasets. We used models of degree 15 as a balance between accurate representation of shape and manageable data dimensionality. *SPHARM* models were then ordinated using a principal components analysis (PCA) of the harmonic coefficients.

To visualize shape change, we generated spherical harmonic representations of shape at -5, -2.5, 0, +2.5, and +5 standard deviations (SDs) along each PC with the *SPHARM* software. The use of +/- 5 SDs ensured that shape differences along axes of otherwise slight shape variation could be identified. To further aid interpretation, we generated contour plots of change relative to the mean shape. We first calculated the Hausdorff distance (Huttenlocher et al., 1993) of the positive and negative standard deviation meshes from the mean shape along that principal component using the ”sampling” filter in `Meshlab` and selecting the ”Distance from the reference mesh” option (Cignoni et al., 1998). Next, we chose the “Show Quality Contour” option under “Render” to produce the contour plot. These plots visualize shape change by gradient, with red hues representing an expansion relative to the reference mesh and blue hues representing a contraction in shape relative to the reference mesh.

### Comparative Analysis

#### Phylogenetic Signal

We assessed phylogenetic signal in our Procrustes and *SPHARM* shape data using K_*mult*_ (Adams, 2014), *the multivariate extension of Blomberg’s K* (Blomberg et al., 2003), as calculated in the `physignal`() function in the `geomorph` library. As a phylogenetic framework we used the maximum clade credibility tree from Slater and Friscia (2019), which was inferred from a supermatrix of nuclear and mitochondrial genes under a relaxed uncorrelated molecular clock and fossilized birth death tree prior. We assessed significance of phylogenetic signal, relative to a null hypothesis of no phylogenetic signal, based on 1000 permutations.

#### Linear Models

To assess the degree to which body mass, locomotor mode, and foot posture predict calcaneal shape in our landmark and *SPHARM* data while accounting for shared evolutionary history, we used Phylogenetic Generalized Least Squares analyses (PGLS) using the `gls`() function from the `nlme` package. For each shape data type, we fit 10 PGLS models that considered each explanatory variable individually, considered pairs of explanatory variables with additive and interactive effects, and all explanatory variables simultaneously with additive effects only. We co-estimated Pagel’s λ by using the `corPagel` correlation structure in the `ape` library (Paradis and Schliep, 2019) to ensure that phylogenetic autocorrelation in the residual error term was appropriately modeled (Revell, 2010). Body mass was transformed logarithmically prior to running the PGLS analyses. We evaluated model support using Akaike information criterion (AIC) values and AIC weights.

## Results

### Principal Components Analysis

Principal components analysis of Procrustes shape variables derived from the landmark data yielded four significant PCs, based on the broken stick method (Frontier, 1976), that explained 31.4, 17.1, 11.0 and 7.4% (total = 66.9%) of shape variation. The first principal component is associated with a continuum from short, broad calcanea with expanded sustentaculae and deep astragalar articular surfaces (negative scores) to long, narrow calcanea with small sustentaculae and shallow astragalar articular surfaces (positive scores). The second principal component describes a continuum from calcanea with long calcaneal heads and an anteroposteriorly long but laterally shallow peroneal tubercle (negative scores), to calcanea with a short calcaneal head and a more anteriorly placed, laterally expanded peroneal tubercle (positive scores). Shape change along PCs 3 and 4 is more difficult to interpret from thin-plate spline deformation grids but involves similar overall patterns of expansion / contraction and elongation / shortening of the calcaneal head relative to the tuber. To determine which axes, if any, correlate with gear ratio, we digitally measured each mesh in Geomagic DesignX. We found a significant positive relationship (*t* = −12.48, *P <* 0.001) between PC1 and gear ratio, accounting for 59% of shape variation. The relationship between Procrustes shape data and gear ratio remained significant but fell to 7% along PC2 (*t* = −2.88, *P <* 0.05) and a relationship was not significant for PCs 3 and 4.

The first 18 principal components of the *SPHARM* coefficients were significant based on the broken stick method. PC1 accounts for approximately 18.6% of shape variation and is interpreted here as describing a gradient from short, stout calcanea at the negative end (mostly represented by members of Ursidae, Mustelidae, and Viverridae) to long, narrow calcanea at the positive end (best represented by Canidae and Felidae). Contour maps illustrate a lateral expansion of the calcaneal tuber and peroneal tubercle and a medial expansion of the sustentaculum (red hues) associated with negative PC1 scores. The positive end of PC1 displays a contraction of the tuber on the lateral side (blue hues) as well as an expansion of the proximal ends of the ectal and sustentacular facets (red hues). The cuboid facet is also markedly different along this axis, with a strong concavity associated with negative PC1 scores and a distinct dorsal lip associated with positive PC1 scores.

PC2 (14.2% of the variance explained) is characterized by the location of the ectal facet (more proximally in negative scores and more distally in positive scores) and, more importantly, the placement of the sustentaculum. The change in the shape of the cuboid facet is similar to that of PC1, where the shape is more concave at one end (positive scores) and more convex at the other (negative scores). The general shape of the calcaneus along PC2, particularly in the tuber, changes from a pronounced dorsal ridge along the tuber with a prominent groove for the Achilles tendon at negative PC2 scores, to a more uniform tuber with a flattened proximal head where the Achilles tendon would attach at positive PC2 scores. The relationship between gear ratio and shape was significant for the first two PCs of SPHARM data (PC1 = 34%, *t* = 7.48, *P <* 0.001; PC2 = 18%, *t* = −4.9, *P <* 0.001), but barely significant for PCs 3 and 4.

### Phylogenetic Signal

The general separation of families that can be identified in the plot of PC1 vs PC2 for Procrustes shape variables translates into a moderate phylogenetic signal that is significantly different from random expectation (*K*_*mult*_ = 0.352, *P <* 0.001). Phylogenetic signal for general shape from the *SPHARM* analyses is slightly lower than for the Procrustes shape data, but significantly greater than zero (*K*_*mult*_ = 0.2479, *P <* 0.001).

### Linear Models

We assessed how the shape of the calcaneus, described by PC scores, is related to locomotion, foot posture, and mass using PGLS analyses for our Procrustes and *SPHARM* data. The best model for Procrustes shape (Table 1, Table S4) is an additive function of locomotion and mass (AIC_*w*_=0.32). However, three other models fall within 2 AIC units of this best model, and therefore we consider them plausible alternatives. All of these models include locomotion as an explanatory variable but have different combinations of additive effects (Table 1). No models including interaction terms were well supported. Effect sizes are large for all locomotor behaviors for the Procrustes shape data, but mass is never significant (Table 1).

**Table 1:** Relative model support (AICw) and coefficient effect sizes for the top four linear models fitted to the Procrustes and SPHARM shape data. Effect sizes are t-statistics, and provided for intercepts and, where applicable, locomotor categories, foot posture, body mass, and interaction terms. Note that the intercept term in models including locomotor mode and/or foot posture is the (composite) effect of arboreal locomotion and/or a digitigrade foot posture on the mean shape.

| Shape Data | Model | $AIC_w$ | Intercept | Locomotor Categories | | | | Posture | Mass | Interaction |
| --- | --- | --- | --- | --- | --- | --- | --- | --- | --- | --- |
|  |  |  |  | **S** | **T** | **F** | **N** |  |  |  |
| Procrustes | locomotion + mass | 0.32 | 3.13 | -3.6 | -4.12 | -3.11 | -2.96 | - | -1.49 | - |
|  | locomotion | 0.29 | 3.14 | -3.66 | -4.43 | -3.34 | -3.46 | - | - | - |
|  | loco + mass + posture | 0.21 | 3.01 | -2.98 | -3.32 | -2.95 | -2.32 | 1.04 | -1.67 | - |
|  | loco + posture | 0.13 | 2.61 | -3.17 | -3.79 | -3.24 | -2.97 | 0.7 | - | - |
| SPHARM | mass + posture | 0.41 | -1.59 | - | - | - | - | -2.14 | 1.98 | - |
|  | mass * posture | 0.25 | -1.87 | - | - | - | - | 0.54 | 2.19 | -1.03 |
|  | posture | 0.16 | 0.11 | - | - | - | - | -1.93 | - | - |
|  | mass | 0.15 | -1.69 | - | - | - | - |  | 1.75 |  |

The best model for our *SPHARM* data was a model with shape as an additive function of mass and foot posture (AIC_*w*_=0.41), with foot posture having the stronger, but not significant, effect (Table 1, Table S4). Three other models fall within ∼ 2 AIC units of this model and include foot posture as a term either by itself (AIC_*w*_ = 0.25) or in an interaction with mass (AIC_*w*_ = 0.16), or include mass as the sole explanatory variable (AIC_*w*_ = 0.15). However, effect sizes are small for all predictors in these models.

## Discussion

A vast toolkit is now available to comparative biologists for quantitatively describing the shapes of complex three-dimensional structures and testing ecological and functional hypotheses about their evolution Adams et al. (2013). However, it is often not clear whether more complete descriptions of shape will yield stronger support for form–function relationships, or whether a few carefully chosen landmarks or measurements might yield comparable support (Gould, 2014). We quantified the shape of the calcaneus in carnivoran mammals using a traditional 3D method based on landmarks (Generalized Procrustes Analysis), as well as a less commonly used approach (*SPHARM*) that quantifies shape variation across the entire surface to assess the relationship between shape and three potential predictors (locomotor mode, body mass, and foot posture) in a phylogenetic context. As expected, phylogenetic regressions suggest that variation in Procrustes shape data is best explained by differences in locomotor mode. We predicted that the *SPHARM* data would show a stronger relationship with locomotor mode compared to the Procrustes shape data, as *SPHARM* provides a more complete description of shape. However, we found allometry and foot posture provided better explanations for overall shape than did locomotor mode, though neither exerted a strong or significant effect. Our results suggest that carefully selected landmarks may be useful for detecting form-function relationships in shape analyses, but that they may over-estimate the effects of ecology on the evolution of anatomical shape in some cases.

Numerous studies have shown that calcaneal shape in mammals is predicted by locomotor mode (Polly, 2008; Panciroli et al., 2017; Polly and MacLeod, 2008; Polly and Sarwar, 2014; Polly, 2020, 2010; Bassarova et al., 2009; Boyer et al., 2013; Jasinski and Wallace, 2014) or aspects of ecology that indirectly influence movement, such as habitat type and vegetation cover (Youlatos, 2003; Polly et al., 2017; Polly and Sarwar, 2014; Polly, 2020; Curran, 2015, 2012). The major axes of shape variation recovered in our Procrustes analyses corroborate previous studies of carnivorans (Polly, 2008; Panciroli et al., 2017; Polly and MacLeod, 2008; Polly and Sarwar, 2014; Polly, 2020, 2010) by showing a continuum from short, broad calcanea, with laterally expanded sustentaculae to long, narrow calcanea with small, proximally placed sustentaculae. Furthermore, we document an associated lengthening of the ectal facet in short/stout calcanea, as found by the 2D landmarking approach of Panciroli et al. (2017). This feature suggests an increase in articular surface area with the astragalus that may promote rotation of the joint (Jenkins Jr and McClearn, 1984; McClearn, 1992; Meldrum et al., 1997; Marsh et al., 2021). It should be noted that these characteristics co-occur most frequently in arboreal, semi-fossorial and natatorial groups such as bears, skunks, mustelids, raccoons, the red panda, and African palm civet (negative PC1 scores, Fig. 3A). Long and narrow calcanea tend to occur in more terrestrial taxa, such as canids, felids, hyaenas, and mongooses (Fig. 3), where the potential for medio-lateral ankle mobility is presumably less biomechanically important for joints that primarily operate in the parasagittal plane.

**Figure 3:**
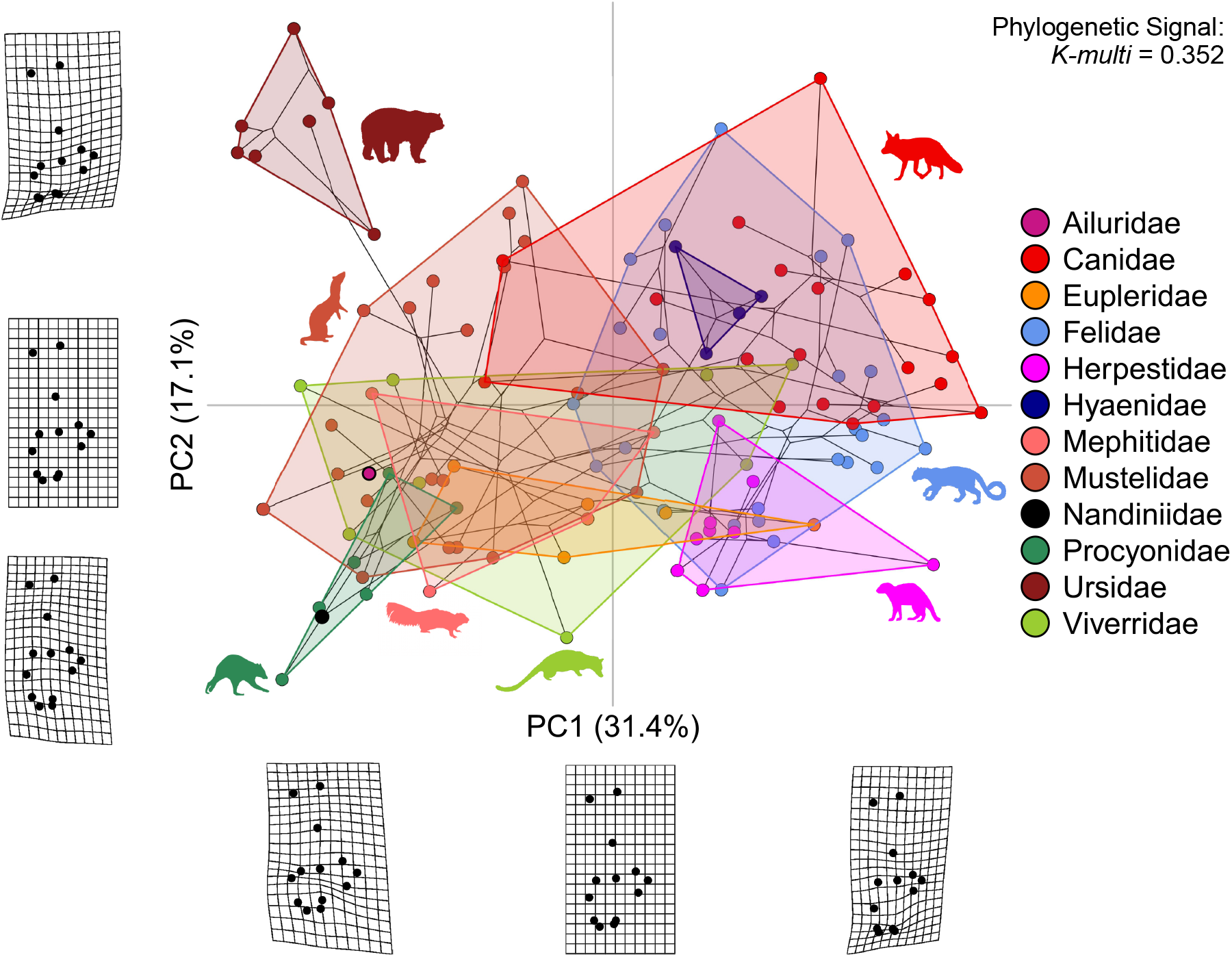
Phylomorphospace for the first two principal components for the Procrustes shape coordinates. The axis values for PC1 were flipped to align with the layout in (Fig. 4). Phylogenetic signal (calculated using Blomberg’s K for multivariate data) is given in the upper right corner. Thin-plate splines illustrate shape changes associated with the extremes, relative to the mean shape (middle) along the two axes. The convex hulls indicate the range for each polytypic carnivoran family.

Changes in calcaneal dimensions also affect gear ratio, which serves as a proxy for hindlimb lever mechanics and foot posture (Polly, 2010). Changes in gear ratio can be observed by examining the placement of the sustentaculum, where a more proximal location increases the effective length of the tuber, increasing the lever arm of the gastrocnemius and soleus muscles, which are important for leaping and running. Thus, high gear ratios are more indicative of digitigrady (Polly, 2010), which tends to be associated with cursorial or terrestrial locomotor strategies (Polly, 2010). Gear ratio accounts for approximately 59% of variation along PC1, which is visually demonstrated by the change in position of the sustentaculum in the thin plate splines (Fig. 3). With all four best-supported linear models for the Procrustes data all including it as a predictor variable, it would seem that calcaneal shape represented by 3D landmarks is most influenced by locomotor mode (Table 1, Table S4). Although models with comparable support include locomotion and body mass as additive effects and a model including locomotion, body mass, and foot posture received comparable support (Table 1), locomotor mode, especially the terrestrial group, had the strongest effect in all of these models effects of foot posture and body mass were small (table 1).

Methods for quantifying shape have evolved over time, with more recent advances providing increasingly accessible ways to more fully describe three-dimensional structures and to more precisely answer questions about form-function relationships. Linear measurements and landmark-based geometric morphometric techniques are practical and computationally efficient, but are not well-suited for characterizing complex 3D features that lack numerous homologous features. Some earlier efforts to characterize variation in 3D shape included an interpolation approach in which a 3D “fishnet” grid of surface points was wrapped around the object, which worked regardless of homologous features being present (Polly, 2008; Polly and MacLeod, 2008). More recently, workers have elaborated on the more traditional landmarking approaches for geometric morphometrics, such as the landmark-free generalized Procrustes surface analysis proposed by Pomidor et al. (2016) which attempts to automate the process of superimposition by employing a refined symmetric iterative closest point approach. Adequately describing a shape with landmarks takes time, and new methods for automating landmarking procedures are particularly useful (Boyer et al., 2015; Felice and Goswami, 2018; Goswami et al., 2019; Bardua et al., 2019), but may still be computationally or labor intensive to implement. We used *SPHARM* because it requires only a small number of homologous landmarks early in the workflow to orient objects to a reference mesh, while still allowing a full description of 3D shape in a robust geometric framework (McPeek et al., 2008; Shen et al., 2009). While we used 13 landmarks for the purpose of comparing our Procrustes analyses to that of Polly et al. (2017), as few as five landmarks were sufficient to orient specimens, corroborating that these methods can be applied to a wide array of complex shapes with minimal guiding landmarks (Curtis and Van Valkenburgh, 2014; Vander Linden et al., 2019). We expected that *SPHARM* would effectively quantify differences in gross calcaneal shape, given the relatively simple geometry of the bone compared to structures such as skulls, and we expected that this would produce greater support for the relationship between shape and locomotor mode, as *SPHARM* uses the entire surface of an object for analysis.

Although PCA of the harmonic coefficients showed general patterns consistent with the long/narrow to short/stout spectrum along PCs 1 and 2, there was more overlap among the carnivoran families and lower phylogenetic signal compared to the Procrustes analysis. Furthermore, although shape change along PC1 could be interpreted as short/stout vs. long/narrow, *SPHARM* more clearly shows that the variation along this continuum is best described as a “robust” to “gracile” gradient. *SPHARM* therefore seems to capture aspects of robustness that cannot be adequately described by the 13 landmarked points alone. This is especially well visualized with contour plots of shape change in the tuber region (Fig. 4). The calcaneal tuber has no recognizable features or homologous landmarks, making it difficult to quantify shape changes in this part of the bone using landmarks. Despite this, *SPHARM* was able to show dramatic changes in the robustness of the tuber. The negative end of PC1 shows a thick tuber with a blunt, well-rounded heel that is characteristic of plantigrade species. In contrast, there is a pronounced groove for the Achilles tendon at the end of a long, narrow tuber in positive PC1 scores, related to digitigrade foot postures. *SPHARM* indicates that there is important variation captured by the calcaneal tuber that has not been quantified before, and is worth investigating further, possibly in the form of cross-sectional dimensions taken at the tuber mid-line as performed by Barr (2020) in bovid calcanea.

**Figure 4:**
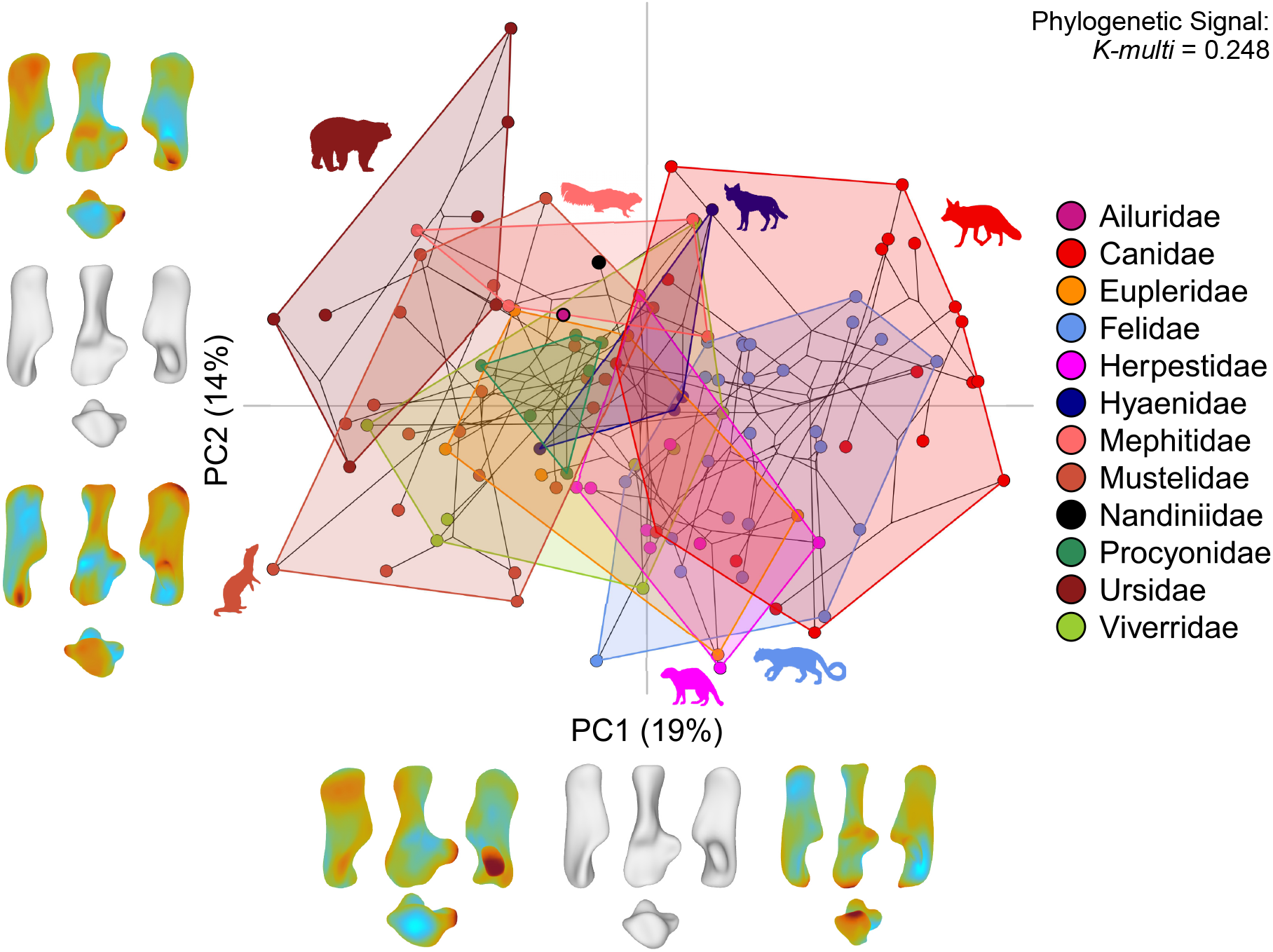
Plot displaying the first two principal components of the *SPHARM* output for general calcaneus shape. Phylogenetic signal (calculated using Blomberg’s K for multivariate data) is given in the upper right corner. *SPHARM* models along each principal component represent shape change along the x axis from the left (−2.5 sd), center (mean) and right (+2.5 sd). *SPHARM* models along the y axis represent shape change along principal component 2 with the top model showing +5 standard deviations from the mean (middle) and the bottom model showing -5 standard deviations from the mean.

Kubo et al. (2019) showed that changes in foot posture are related to the rate of evolution of body size, in that the adoption of more upright foot postures is associated with increases in body mass. The idea that mammalian calcaneal shape may have a strong allometric component is also indicated by several studies that have predicted mammalian body masses from calcaneal measurements (Yapuncich et al., 2015; Tsubamoto, 2019). Consistent with these ideas, but in contrast with the results from analysis of Procrustes shape data, linear modeling suggests that the main contributors to carnivoran calcaneal shape change, as described by spherical harmonics, are body mass and foot posture, though their effects are weak overall (Table 1, Table S4). It is unclear exactly why shape described by GPA and *SPHARM* yield conflicting support for drivers of calcaneal shape evolution, but an obvious explanation is that Polly et al.’s (2017) landmarks focus on shape change related to gear ratio (refer to landmarks 1–3, 6, and 12–13 and equation B/A in Fig. 1), as well as aspects of the astragalo-calcaneal articulation, such as the length and position of the ectal facet (landmarks 9–11), and the shape of the sustentacular facet (landmarks 6–8), which must also provide a strong locomotor signal (Panciroli et al., 2017; Ginot et al., 2016). These features are also clearly captured by *SPHARM* (Fig. 4), but this method’s ability to describe other aspects of shape that cannot be adequately represented by landmarks results in a more complete description of shape that is not only focused on articular surfaces that play an important role in locomotion. The fact that our “best” linear models for the *SPHARM* data all yield low and insignificant effects for foot posture, body mass, and locomotor behavior suggest that patterns of shape variation in this dataset are, in fact, more complex that these simple models can predict. Regardless of whether or not foot posture and allometry together provide a better overall explanation for patterns of shape variation in carnivoran calcanea than does locomotor behavior, it is apparent that the signal emerging from our Procrustes data is not supported by analysis of the shape of the entire bone and that generalizing patterns of articular surface shape evolution to the calcaneus as a whole are premature.

While it is clear that the relative influence of the three predictors on macroevolutionary shape change differ based on the method used to quantify shape, it is possible that our groupings for foot posture and locomotor mode are too coarse to extract meaningful ecological signal, regardless of the type of morphometric data. Limiting foot posture categories to only plantigrade and digitigrade may be too simple of a strategy, though the degree to which some carnivorans engage in semi-digitigrade behaviors is unclear and would benefit from more rigorous assessment. More importantly, our strategy for assigning locomotor groups, though based on many studies predating this one (Van Valkenburgh, 1985, 1987; Polly, 2008; Polly and MacLeod, 2008; Polly, 2010; Kilbourne and Hoffman, 2013, 2015; Kilbourne, 2017; Samuels et al., 2013; Panciroli et al., 2017) and a standard mammalian locomotor categorization (Eisenberg, 1981), is likely inadequate to capture the true diversity of behaviors employed by carnivorans or other mammal groups. For example, “terrestrial” here encompasses taxa as diverse as large cats, bears, canids, and hyaenas, and therefore may overlook functionally different ways of being terrestrial. For carnivorans specifically, reconsidering the way we diagnose locomotor behavior may be further aided by incorporating predatory habits, as felids often leap and employ ambush tactics (Polly, 2020) while many canids engage in long-distance pursuit predation, each of which likely loads the hindlimb in different ways. Similarly, scansorial and arboreal behaviors may blend into one another, resulting in challenges to simple classification. Nations et al. 2019 calculated a “climb index” based on trapping records for 20 murine rodents which provided a more quantitative and natural approach for diagnosing the degree of terrestrial versus arboreal behavior. In order to avoid the arbitrary and imprecise categorization of mammals into dietary groups, **?** used an ordinal ranking system to quantify the relative importance of food items in the diets of mammals. Development of a similar coding scheme for mammalian substrate use could serve as a model for incorporating more behavior-based ways of attributing locomotor mode to mammals and other vertebrates, yielding novel insights into the relationships between osteological form and locomotor behavior.

The evolutionary processes at play within carnivorans regarding the relationships between calcaneal shape, foot posture and body mass could be further explored in the future by making use of the excellent fossil record that this clade possesses. Ursids present a particularly interesting case study. As in previous work (Polly, 2010; Panciroli et al., 2017; Polly et al., 2017), we found that bears have rather distinct calcaneal shapes relative to other extant carnivorans (Figures 3,4). Despite being the largest extant terrestrial carnivorans and including the largest extinct terrestrial carnivorans to have ever lived (Christiansen, 1999; Soibelzon and Schubert, 2011), some extinct ursids achieved body sizes more comparable to small extant carnivorans (Finarelli and Flynn, 2006) and may have had more arboreal habits. It is therefore possible that the “ambulatory” locomotor mode of extant ursids is a constraint, rather than an adaptation, that results from a foot posture and morphology that is more suited to arboreal behaviors at small body size than terrestrial locomotion at large sizes. We also acknowledge that, while we have the largest and most diverse sample of carnivorans in a study of three-dimensional calcaneal shape, this sample is still not large enough to rigorously test remaining questions about shape diversity using data-hungry comparative methods. Although the fossil record provides one way to supplement our sample, the remaining 4,000+ extant mammalian species (Burgin et al., 2018) span a range of ecologies, morphologies, and behaviors that greatly exceed that seen in Carnivora (e.g., Boyer et al., 2013; Dunn and Avery, 2021), and may provide further insight into how the change in calcaneal shape corresponds to aspects of ecology, behavior, and phylogeny. The landmark-deficient nature of the calcaenus presents barriers to robust analysis using traditional morphometric approaches, but *SPHARM* ‘s ability to describe shape in a landmark-free framework makes it an ideal tool for future work in this area.

## Authorship Contributions

GJS conceived the study; ANW,RN, and GJS scanned specimens, ANW, RN, and RH processed scan data and prepared models for analysis, ANW, RN, and GJS performed analyses, ANW and GJS wrote the manuscript.

## Acknowledgements

We thank Adam Ferguson for providing access to specimens. Ken Angielcyzk, Caroline Abbott, Jacqueline Lungmus, Isaac Magallanes, Jason Pardo, Tristan Reinecke, Daniel Rhoda, Stephanie Smith, Sarah Saxton Strassberg, Adrienne Stroup, Chloe Nash, Zhe-Xi Luo, Jonathan Nations provided comments on earlier drafts of the manuscript.

