## Supplementary Data for "Choice of 3D morphometric method leads to diverging interpretations of form–function relationships in the carnivoran calcaneus"

Version dated: May 16, 2022

running title: Morphometrics and carnivoran calcaneal shape

[3] *School of GeoSciences, University of Edinburgh, U.K.*

### Supplementary Tables

Table S1: Table with the PANtheria Jones et al. (2009) average adult body mass values for carnivoran taxa. If information was not available we used data from its closest relative.

| Species | Mass (g) | Notes |
|---|---|---|
| Acinonyx jubatus | 50577.92 | |
| Ailuropoda melanoleuca | 117999.99 | |
| Ailurus fulgens | 5170.08 | |
| Aonyx cinereus | 3527.59 | |
| Arctictis binturong | 12999.99 | |
| Arctogalidia trivirgata | 2323.79 | |
| Arctonyx albogularis | 8166.52 | used A. collaris |
| Atelocynus microtis | 8363.22 | |
| Atilax paludinosus | 3600.16 | |
| Bassaricyon alleni | 1250 | used B. gabbii |
| Bassariscus astutus | 1010.37 | |
| Bdeogale nigripes | 2623.01 | |
| Canis aureus | 9658.7 | |
| Canis latrans | 11989.1 | |
| Canis lupus | 31756.51 | |
| Canis mesomelas | 8247.3 | |
| Caracal aurata | 11277.17 | |
| Caracal caracal | 11964.38 | |
| Catopuma temminckii | 7726.46 | |
| Cerdocyon thous | 5741.66 | |
| Chrysocyon brachyurus | 23325 | |
| Civettictis civetta | 12075.58 | |
| Conepatus leuconotus | 3293.91 | |
| Crocuta crocuta | 63369.98 | |
| Cryptoprocta ferox | 9500 | |
| Cuon alpinus | 15800 | |
| Eira barbara | 4134.99 | |
| Enhydra lutris | 27410.93 | |
| Eupleres goudotii | 2763.34 | |
| Felis chaus | 7157.99 | |
| Felis margarita | 2823.36 | |
| Felis silvestris | 4573.08 | |
| Fossa fossana | 1853.98 | |
| Galictis vittata | 1000 | used G. cuja |
| Galidia elegans | 810 | |
| Galidictis fasciata | 549.99 | |
| Gulo gulo | 12792.49 | |
| Helarctos malayanus | 57075.78 | |

| species | Mass (g) | Notes |
|---|---|---|
| Helogale parvula | 281.83 | |
| Hemigalus derbyanus | 1262.7 | |
| Herpailurus yagouaroundi | 6875 | |
| Herpestes ichneumon | 2980.02 | |
| Herpestes sanguineus | 543.83 | |
| Hyaena hyaena | 35070.51 | |
| Hydrictis maculicollis | 4180.53 | |
| Ichneumia albicauda | 3628.4 | |
| Ictonyx striatus | 811.02 | |
| Leopardus colocola | 8133.78 | used L. jacobitus |
| Leopardus pardalis | 11880 | |
| Leopardus tigrinus | 8133.78 | used L. jacobitus |
| Leopardus wiedii | 3270.81 | |
| Leptailurus serval | 11999.96 | |
| Lontra canadensis | 8087.42 | |
| Lontra longicaudis | 6554.97 | |
| Lupulella adustus | 10392.49 | |
| Lutra lutra | 8868.69 | |
| Lycalopex griseus | 4542.67 | used L. gymnocercus |
| Lycaon pictus | 21999.99 | |
| Lynx canadensis | 9682.82 | |
| Lynx rufus | 6374.47 | |
| Martes americana | 873.69 | |
| Meles meles | 11884.03 | |
| Mellivora capensis | 8999.99 | |
| Melursus ursinus | 110500 | used U. americanus |
| Mephitis mephitis | 2399.99 | |
| Mungos mungo | 1260 | |
| Mungotictis decemlineata | 657.03 | |
| Mustela frenata | 190.03 | |
| Mustela putorius | 975.55 | |
| Mustela vison | 190.03 | used M. frenata |
| Mydaus javanensis | 2500 | |
| Nandinia binotata | 2167.2 | |
| Nasua nasua | 3775.5 | |
| Neofelis nebulosa | 14945.05 | |
| Nyctereutes procyonoides | 4214.99 | |
| Otocolobus manul | 3050 | |
| Otocyon megalotis | 4098.12 | |
| Paguma larvata | 4300 | |

| species | Mass (g) | Notes |
|---|---|---|
| Panthera leo | 158623.93 | |
| Panthera onca | 83943.09 | |
| Panthera pardus | 52399.99 | |
| Panthera tigris | 161914.66 | |
| Panthera uncia | 32500 | |
| Paradoxurus hermaphroditus | 3200 | |
| Parahyaena brunnea | 42977.93 | |
| Pardofelis marmorata | 2826.68 | |
| Pekania pennanti | 3750 | |
| Poecilogale albinucha | 308.16 | |
| Potos flavus | 2441.81 | |
| Prionailurus bengalensis | 2780.97 | |
| Procyon cancrivorus | 6931.72 | |
| Procyon lotor | 6373.72 | |
| Proteles cristata | 8139.39 | |
| Pteronura brasiliensis | 26000 | |
| Puma concolor | 53954.05 | |
| Speothos venaticus | 6324.54 | |
| Spilogale putorius | 566.49 | |
| Suricata suricatta | 729.99 | |
| Taxidea taxus | 7842.15 | |
| Tremarctos ornatus | 123176.97 | |
| Urocyon cinereoargenteus | 3833.71 | |
| Ursus americanus | 110500 | |
| Ursus arctos | 196287.5 | |
| Ursus maritimus | 371703.81 | |
| Urva javanica | 750 | |
| Viverra zibetha | 9148.77 | |
| Viverricula indica | 2918.88 | |
| Vulpes lagopus | 3584.37 | |
| Vulpes velox | 2088 | |
| Vulpes vulpes | 4820.36 | |
| Vulpes zerda | 1317.13 | |

Table S2: Table with species information and references for locomotor mode. We chose locomotor mode based on a consensus from what previous papers had assigned species. The main papers we used constitute columns 3-12 and are as follows: P&MO8,Polly and MacLeod (2008); P08,Polly (2008); Pan17, Panciroli et al. (2017); VV87,Van Valkenburgh (1987); S13, Samuels et al. (2013); VV85, Van Valkenburgh (1985); K13, Kilbourne and Hoffman (2013); K15, Kilbourne and Hoffman (2015); K17, Kilbourne (2017). Other references were used if there was one or no previous assignments from other studies. Abbreviations are as follows: T = terrestrial, A = arboreal, S = scansorial, F = fossorial/semifossorial, N = natatorial/semi-aquatic, C = cursorial, G = generalized, L = ambulatory. We considered cursorial, generalized, and ambulatory as part of the terrestrial category (see main text). We made several substitutions based on available data: *Bassaricyon alleni* (used *B. gabbii* in Polly (2008); Polly and MacLeod (2008); Polly (2010)), *Arctonyx albogularis* (used *A. collaris* in Panciroli et al. (2017); Polly (2010); Van Valkenburgh (1987); Samuels et al. (2013); Kilbourne (2017)), *Conepatus leuconotus* (used *C. chinga* in Panciroli et al. (2017)), *Lycalopex griseus* (used *L. gymnocerus* in Panciroli et al. (2017) and *Lycalopex* sp. in Samuels et al. (2013)), *Nasua nasua* (used *N. narica* in Van Valkenburgh (1987)), and *Bdeogale nigripes* (used *B. crassicauda* and *B. jacksoni* in Samuels et al. (2013)).

| Species | Mode | P&M08 | P08 | Pan17 | P10 | VV87 | S13 | VV85 | K13 | K15 | K17 | other |
|---|---|---|---|---|---|---|---|---|---|---|---|---|
| Acinonyx jubatus | T | | | C | S | S | C | S | C | C | | |
| Ailuropoda melanoleuca | T | | | L | S | | T | | | | | Nowak and Walker (1999) |
| Ailurus fulgens | A | A | A | A | A | | A | | | | | Reid et al. (1991) |
| Aonyx cinereus | N | | | | | | N | | | | N | |
| Arctictis binturong | A | | | A | A | A | A | A | | | | |
| Arctogalidia trivirgata | A | | | A | | | | | | | | Nowak and Walker (1999); Barca et al. (2016); Moore (2011); Willcox et al. (2012) |
| Arctonyx albogularis | F | | | F | F | F | F | | | | F | |
| Atelocynus microtis | T | | | | T | | T | | | | | |
| Atilax paludinosus | N | | | N | N | | N | | | | | |
| Bassaricyon alleni | A | A | A | A | A | | | | | | | |
| Bassariscus astutus | S | | | A | T | S | A | | | | | Nowak and Walker (1999); Frey and Godding (2013); Poglayen-Neuwall and Toweill (1988) |
| Bdeogale nigripes | T | | | | T | | T | | | | | |
| Canis aureus | T | | | T | T | T | | T | | | | |
| Canis latrans | T | | | C | T | T | C | T | C | C | | |
| Canis lupus | T | | | C | T | T | C | T | C | C | | |
| Canis mesomelas | T | | | C | T | T | C | T | | | | |
| Caracal aurata | T | | | | S | | | | | | | Nowak and Walker (1999); Bahaa-el din (2015) |
| Caracal caracal | S | | | S | S | S | | S | | C | | |
| Catopuma temminckii | S | | | | S | S | | S | | | | |

| Species | Mode | P&M08 | P08 | Pan17 | P10 | VV87 | S13 | VV85 | K13 | K15 | K17 | other |
|---|---|---|---|---|---|---|---|---|---|---|---|---|
| Cerdocyon thous | T | | | | T | | T | T | | | | |
| Chrysocyon brachyurus | T | | | T | T | | T | T | | | | |
| Civettictis civetta | T | | | | S | | T | T | | | | Nowak and Walker (1999) |
| Conepatus leuconotus | F | | | F | T | | F | | | | | Nowak and Walker (1999); Dragoo and Sheffield (2009); Brashear et al. (2010) |
| Crocuta crocuta | T | T | T | C | T | T | C | T | | | | |
| Cryptoprocta ferox | A | | | A | S | | A | | | | | Nowak and Walker (1999) |
| Cuon alpinus | T | | | C | T | T | C | T | | | | |
| Eira barbara | S | | | A | S | S | A | | | | S | Nowak and Walker (1999) |
| Enhydra lutris | N | | | | N | | N | | | | N | |
| Eupleres goudotii | T | | | T | | | | | | | | Nowak and Walker (1999) |
| Felis chaus | S | | | | S | | | | | | | |
| Felis margarita | S | | | | S | | | | | | | |
| Felis silvestris | S | | | | | | S | | | | | Nowak and Walker (1999) |
| Fossa fossana | T | | | S | T | | T | | | | | Nowak and Walker (1999) |
| Galictis vittata | F | | | | F | T | T | | | | G | |
| Galidia elegans | S | | | | | | S | | | | | Nowak and Walker (1999) |
| Galidictis fasciata | T | | | | | | T | | | | | |
| Gulo gulo | T | | | T | F | S | T | S | | | G | Nowak and Walker (1999) |
| Helarctos malayanus | S | | | L | S | S | S | S | | | | |
| Helogale parvula | T | | | | | | | | | | | Nowak and Walker (1999); Taylor (1976) |
| Hemigalus derbyanus | S | | | | S | | S | | | | | |
| Herpailurus yagouaroundi | S | | | S | S | | | | | | | |
| Herpestes ichneumon | T | | | T | T | | T | | | | | |
| Herpestes sanguineus | T | | | | | | T | | | | | Nowak and Walker (1999) |
| Hyaena hyaena | T | | | T | T | T | C | T | | | | |
| Hydrictis maculicollis | N | | | | | | | | | | N | Nowak and Walker (1999) |
| Ichneumia albicauda | T | | | T | T | T | T | | | | | |
| Ictonyx striatus | F | | | | S | F | F | | | | F | Nowak and Walker (1999) |
| Leopardus colocola | S | | | S | | | | | | | | Nowak and Walker (1999) |
| Leopardus pardalis | S | | | S | S | S | | S | | | | |

| Species | Mode | P&M08 | P08 | Pan17 | P10 | VV87 | S13 | VV85 | K13 | K15 | K17 | other |
|---|---|---|---|---|---|---|---|---|---|---|---|---|
| Leopardus tigrinus | S | | | | S | | | | | | | |
| Leopardus wiedii | A | | | A | S | A | A | | | | | |
| Leptailurus serval | T | T | T | T | S | | T | S | | | | |
| Lontra canadensis | N | | | N | N | | N | | N | N | N | |
| Lontra longicaudis | N | | | | N | | | | | | N | |
| Lupulella adustus | T | | | | T | | C | T | | | | |
| Lutra lutra | N | N | N | N | | | N | | | | N | |
| Lycalopex griseus | T | | | C | T | | C | | | | | |
| Lycaon pictus | T | | | C | T | T | C | T | C | C | | |
| Lynx canadensis | S | | | T | S | S | T | S | | | | Nowak and Walker (1999) |
| Lynx rufus | S | S | S | S | S | | S | S | C | C | | |
| Martes americana | S | | | S | S | | S | | | | S | |
| Meles meles | F | F | F | F | F | F | F | F | | | F | |
| Mellivora capensis | F | | | F | F | F | F | F | F | F | F | |
| Melursus ursinus | S | | | L | S | S | T | S | | | | |
| Mephitis mephitis | F | | | F | T | F | F | | G | G | | |
| Mungos mungo | T | | | | | | T | | | | | Nowak and Walker (1999) |
| Mungotictis decemlineata | S | | | | | | | | | | | Nowak and Walker (1999) |
| Mustela frenata | T | | | | T | | T | | | | G | |
| Mustela putorius | T | T | T | | | | T | | | | G | |
| Mustela vison | N | | | | T | | N | | | N | N | |
| Mydaus javanensis | F | | | | | | | | | | | Fabre et al. (2015) |
| Nandinia binotata | A | | | A | S | | A | | | | | Nowak and Walker (1999) |
| Nasua nasua | S | | | S | S | S | | | | | | |
| Neofelis nebulosa | A | | | A | S | A | A | A | | | | Nowak and Walker (1999) |
| Nyctereutes procyonoides | T | | | T | T | T | T | | | | | |
| Otocolobus manul | T | | | T | S | | T | | | | | |
| Otocyon megalotis | T | | | | T | | | | C | C | | Nowak and Walker (1999) |
| Paguma larvata | A | | | A | | A | A | | | | | |
| Panthera leo | T | | | T | T | S | T | S | | | | |

| Species | Mode | P&M08 | P08 | Pan17 | P10 | VV87 | S13 | VV85 | K13 | K15 | K17 | other |
|---|---|---|---|---|---|---|---|---|---|---|---|---|
| Panthera onca | S | | | S | S | S | | S | | | | |
| Panthera pardus | S | | | S | S | S | | S | | | | |
| Panthera tigris | T | | | T | T | S | | S | | | | |
| Panthera uncia | S | | | S | S | | S | S | | | | |
| Paradoxurus hermaphroditus | A | | A | A | A | A | A | | | | | |
| Parahyaena brunnea | T | | | T | T | | C | | | | | |
| Pardofelis marmorata | S | | | | S | | A | | | | | Nowak and Walker (1999) |
| Pekania pennanti | S | | | | S | S | S | | S | S | S | |
| Poecilogale albinucha | T | | | | T | | | | | | F | Nowak and Walker (1999) |
| Potos flavus | A | | | A | A | A | A | | | | | |
| Prionailurus bengalensis | S | | | S | S | | | | | | | |
| Procyon cancrivorus | S | | | | S | | S | S | | | | |
| Procyon lotor | S | | | S | S | S | S | | S | S | | |
| Proteles cristata | T | | | T | T | | T | | | | | |
| Pteronura brasiliensis | N | | | N | | | | | | | N | |
| Puma concolor | S | | | S | S | S | S | S | C | C | | |
| Speothos venaticus | T | | | | T | T | | T | | | | |
| Spilogale putorius | F | | | | T | | F | | | | | Dumont et al. (2016) |
| Suricata suricatta | T | | | | T | | F | | | | | |
| Taxidea taxus | F | | | F | F | F | F | F | F | F | F | |
| Tremarctos ornatus | S | | | L | S | S | S | | | | | |
| Urocyon cinereoargenteus | S | | | S | S | S | S | | C | C | | |
| Ursus americanus | S | | | L | S | S | S | S | | | | |
| Ursus arctos | T | | | L | S | T | T | T | | | | |
| Ursus maritimus | T | | | L | T | | N | | | | | |
| Urva javanica | T | | | | T | | | | | | | Nowak and Walker (1999) |
| Viverra zibetha | T | | | T | S | | T | T | | | | |
| Viverricula indica | T | | | T | S | | T | | | | | Nowak and Walker (1999) |
| Vulpes lagopus | T | | | T | T | | T | | | | | |
| Vulpes velox | T | | | T | T | | | | | | | |

| Species | Mode | P&M08 | P08 | Pan17 | P10 | VV87 | S13 | VV85 | K13 | K15 | K17 | other |
|---|---|---|---|---|---|---|---|---|---|---|---|---|
| Vulpes vulpes | T | | | C | T | T | C | T | C | C | | |
| Vulpes zerda | T | | | C | T | | C | | | | | |

Table S3: Table with species information and references for foot posture. Abbreviations are as follows: P = plantigrade, S = semi-digitigrade, D = digitigrade. We used a consensus approach when assigning foot posture for the species in our analyses based on other studies. Main resources are in columns 3-6, with supplementary references in column 7. Studies are: P&M08,Polly and MacLeod (2008); P08, Polly (2008);P10,Polly (2010);C97,Carrano (1997). We made several substitutions based on available data: *Bassaricyon alleni* (used *B. gabbii* in Polly (2008); Polly and MacLeod (2008)), and *Arctonyx albogularis* (used *A. collaris* in Polly (2010)). Asterisks in the final column indicate that the posture for the closest relative in our dataset was used if there was no information found in the literature: *Aonyx cinereus* (used *Lutra lutra*), *Helogale parvula* (used *Mungos mungo*), *Hydrictis maculicollis* (used *Lutra lutra*), *Mungotictis decemlineata* (used *Galidictis fasciata*), *Poecilogale albinucha* (used *Ictonyx striatus*), *Leopardus colocola* (used I).

| Species | Posture | P&M08 | P08 | P10 | C97 | other |
|---|---|---|---|---|---|---|
| Acinonyx jubatus | D | | | D | D | |
| Ailuropoda melanoleuca | P | | | P | | |
| Ailurus fulgens | P | P | P | P | P | |
| Aonyx cinereus | D | | | | | * |
| Arctictis binturong | P | | | S | P | Taylor (1988) |
| Arctogalidia trivirgata | P | | | | | Nowak and Walker (1999) |
| Arctonyx albogularis | D | | | S | | |
| Atelocynus microtis | D | | | D | | |
| Atilax paludinosus | D | | | S | | |
| Bassaricyon alleni | P | P | P | P | | |
| Bassariscus astutus | D | | | S | | |
| Bdeogale nigripes | D | | | D | | |
| Canis aureus | D | | | D | | |
| Canis latrans | D | | | D | | |
| Canis lupus | D | | | D | D | |
| Canis mesomelas | D | | | D | | |
| Caracal aurata | D | | | D | | |
| Caracal caracal | D | | | D | | |
| Catopuma temminckii | D | | | D | | |
| Cerdocyon thous | D | | | D | | |
| Chrysocyon brachyurus | D | | | D | D | |
| Civettictis civetta | D | | | D | D | |
| Conepatus leuconotus | D | | | S | | |
| Crocuta crocuta | D | D | D | D | D | |
| Cryptoprocta ferox | P | | | S | P | Nowak and Walker (1999); Köhncke and Leonhardt (1986) |
| Cuon alpinus | D | | | D | D | |
| Eira barbara | P | | | P | | |
| Enhydra lutris | P | | | P | | |
| Eupleres goudotii | D | | | | | Taylor (1988) |
| Felis chaus | D | | | D | | |

| Species | Posture | P&M08 | P08 | P10 | C97 | other |
|---|---|---|---|---|---|---|
| Felis margarita | D | | | D | | |
| Felis silvestris | D | | | | | Ginsburg (1961) |
| Fossa fossana | D | | | D | D | |
| Galictis vittata | P | | | P | P | |
| Galidia elegans | D | | | | | Taylor (1988) |
| Galidictis fasciata | D | | | | | Taylor (1988) |
| Gulo gulo | P | | | S | P | Copeland and Kucera (1997) |
| Helarctos malayanus | P | | | P | | |
| Helogale parvula | D | | | | | * |
| Hemigalus derbyanus | D | | | S | D | |
| Herpailurus yagouaroundi | D | | | D | | |
| Herpestes ichneumon | D | | | S | | |
| Herpestes sanguineus | D | | | | | Kingdon (1977) |
| Hyaena hyaena | D | | | D | | |
| Hydrictis maculicollis | D | | | | | * |
| Ichneumia albicauda | D | | | D | | |
| Ictonyx striatus | P | | | P | | |
| Leopardus colocola | D | | | | | * |
| Leopardus pardalis | D | | | D | | |
| Leopardus tigrinus | D | | | D | | |
| Leopardus wiedii | D | | | D | | |
| Leptailurus serval | D | D | D | D | | |
| Lontra canadensis | D | | | S | | |
| Lontra longicaudis | D | | | S | | |
| Lupulella adustus | D | | | D | | |
| Lutra lutra | D | S | S | | | |
| Lycalopex griseus | D | | | D | | |
| Lycaon pictus | D | | | D | D | |
| Lynx canadensis | D | | | D | | |
| Lynx rufus | D | D | D | D | D | |
| Martes americana | P | | | P | | |
| Meles meles | P | S | S | S | P | Ginsburg (1961) |
| Mellivora capensis | P | | | S | P | Ginsburg (1961) |
| Melursus ursinus | P | | | P | P | |
| Mephitis mephitis | P | | | P | P | |
| Mungos mungo | D | | | | P | Kingdon (1977) |
| Mungotictis decemlineata | D | | | | | * |
| Mustela frenata | D | | | S | | |
| Mustela putorius | D | S | S | | | |
| Mustela vison | D | | | | | * |

| Species | Posture | P&M08 | P08 | P10 | C97 | other |
|---|---|---|---|---|---|---|
| Mydaus javanensis | P | | | | | Ginsburg (1961) |
| Nandinia binotata | P | | | P | | |
| Nasua nasua | P | | | P | P | |
| Neofelis nebulosa | D | | | D | D | |
| Nyctereutes procyonoides | D | | | D | D | |
| Otocolobus manul | D | | | D | | |
| Otocyon megalotis | D | | | D | | |
| Paguma larvata | P | | | | | Taylor (1988) |
| Panthera leo | D | | | D | | |
| Panthera onca | D | | | D | | |
| Panthera pardus | D | | | D | | |
| Panthera tigris | D | | | D | D | |
| Panthera uncia | D | | | D | | |
| Paradoxurus hermaphroditus | P | | S | S | P | Taylor (1988) |
| Parahyaena brunnea | D | | | D | D | |
| Pardofelis marmorata | D | | | D | | |
| Pekania pennanti | P | | | P | | |
| Poecilogale albinucha | P | | | | | * |
| Potos flavus | P | | | P | | |
| Prionailurus bengalensis | D | | | D | | |
| Procyon cancrivorus | D | | | S | | |
| Procyon lotor | P | | | S | P | "Jenkins and Camazine (1977); Lotze and Anderson (1979)" |
| Proteles cristata | D | | | D | | |
| Pteronura brasiliensis | P | | | | | Noonan et al. (2017) |
| Puma concolor | D | | | D | D | |
| Speothos venaticus | D | | | D | | |
| Spilogale putorius | P | | | P | | |
| Suricata suricatta | D | | | S | | |
| Taxidea taxus | P | | | P | P | |
| Tremarctos ornatus | P | | | P | P | |
| Urocyon cinereoargenteus | D | | | D | D | |
| Ursus americanus | P | | | P | | |
| Ursus arctos | P | | | P | P | |
| Ursus maritimus | P | | | P | | |
| Urva javanica | D | | | S | | |
| Viverra zibetha | D | | | D | | |
| Viverricula indica | D | | | D | | |
| Vulpes lagopus | D | | | D | | |
| Vulpes velox | D | | | D | | |
| Vulpes vulpes | D | | | D | D | |

| Species | Posture | P&M08 | P08 | P10 | C97 | other |
|---|---|---|---|---|---|---|
| Vulpes zerda | D | | | D | | |

Table S4: AIC scores, differences from the best model (ΔAIC), relative model support (AICw) and phylogenentic signal (Pagel's $\lambda$ for phylogenetic generalized least squares models fitted to Procrustes and (*SPHARM*) shape data.

| Model | Procrustes | | | | SPHARM | | | |
|---|---|---|---|---|---|---|---|---|
| | AIC | ΔAIC | AICw | $\lambda$ | AIC | ΔAIC | AICw | $\lambda$ |
| shape ∼ locomotion | -429.03 | 0.22 | 0.29 | 0.81 | -1200.66 | 7.36 | 0.01 | 0.69 |
| shape ∼ locomotion + foot posture | -427.52 | 1.74 | 0.13 | 0.79 | -1199.7 | 8.32 | < 0.01 | 0.66 |
| shape ∼ locomotion * foot posture | -420.88 | 8.38 | <0.01 | 0.78 | -1194.58 | 13.44 | <0.01 | 0.68 |
| shape ∼ locomotion + mass | -429.26 | 0 | 0.32 | 0.83 | -1200.86 | 7.16 | 0.01 | 0.71 |
| shape ∼ locomotion * mass | -421.93 | 7.32 | <0.01 | 0.83 | -1195.03 | 12.99 | <0.01 | 0.71 |
| shape ∼ mass | -421.33 | 7.92 | <0.01 | 0.91 | -1205.94 | 2.08 | 0.15 | 0.75 |
| shape ∼ mass + foot posture | -423.78 | 5.47 | 0.02 | 0.87 | -1208.02 | 0 | 0.41 | 0.66 |
| shape ∼ mass * foot posture | -422.132 | 7.12 | <0.01 | 0.86 | -1206.99 | 1 | 0.25 | 0.61 |
| shape ∼ foot posture | -419.792 | 9.46 | <0.01 | 0.84 | -1206.08 | 1.94 | 0.16 | 0.64 |
| shape ∼ locomotion + mass + foot | -428.361 | 0.89 | 0.21 | 0.82 | -1200.94 | 7.08 | 0.01 | 0.67 |

*
